# Emergence of a Dengue virus serotype 2 causing the largest ever dengue epidemic in Sri Lanka

**DOI:** 10.1101/329318

**Authors:** Ananda Wijewickrama, Samitha Fernando, Geethal S. Bandara Jayerathne, Pathum Asela Perera, S.A. Abeynaike, Laksiri Gomes, Chandima Jeewandara, Rashmi Tippalagama, Desha Dilani, Anishgoby Rajendran, Ranmalie Abeyesekere, Suraj Goonawardhana, Aruna Dharshan De Silva, Gathsaurie Neelika Malavige

**Author notes:** Current Address: General Sir John Kotelawala Défense University, Ratmalana, Sri Lanka. These senior authors contributed equally to this article (Ananda Wijewickrama, Aruna Dharshan De Silva, Gathsaurie Neelika Malavige). Correspondence should be addressed to: Prof. Gathsaurie Neelika Malavige DPhil, FRCP, FRCPath Centre for Dengue Research, Faculty of Medical Sciences, University of Sri Jayawardanapura, Sri Lanka.

## Abstract

**Background:** Sri Lanka experienced the largest ever dengue outbreak in year 2017, which coincided with the shift of the predominant circulating dengue virus (DENV) 1 to DENV2 after 9 years. As it was felt that more patients appeared to develop complications and severe dengue, we compared clinical features of patients with acute dengue, with the previous circulating serotype (DENV1) and also sequenced the new virus, to determine the lineage of the virus.

**Methodology/Principal findings:** We studied the clinical and laboratory features of 172 adult patients with acute DENV1 (n=79) and DENV2 (n=93) infection. 65 (82.3%) of those with DENV1 and 86 (92.4%) of those with DENV2 were experiencing a secondary infection. The risk of developing dengue haemorrhagic fever (DHF) was significantly higher (=0.005, odds ratio=2.5) in those infected with DENV2 (54.8%) when compared to DENV1(32.9%), even though similar proportions of patients had a secondary dengue infection. Patients with DENV2 infection developed leakage significantly earlier (p<0.0001, median= 3, days) when compared to those with a DENV1 infection (median 5 days) and were more likely to develop significant bleeding and to require blood transfusions. Furthermore, patients with DENV2 were more likely to have significantly lower platelet counts during day 3, 4 and 5 since onset of illness.

Whole genome sequencing showed that these DENV-2 isolates belonged to a cosmopolitan strain and was genetically more distant than the DENV-2 strains that circulated from 1981 to 2004 in Sri Lanka.

**Conclusions/significance:** Since this DENV2 strain appears to cause more severe forms of clinical disease, it would be important to determine variations in the virus genome or other factors that could have contributed to severe disease.

**Author summary:** Sri Lanka experienced the largest ever dengue outbreak in year 2017, which coincided with the shift of the predominant circulating dengue virus (DENV) 1 to DENV2 after 9 years. We studied the clinical and laboratory features of 172 adult patients with acute DENV1 (n=79) and DENV2 (n=93) infection. The risk of developing dengue haemorrhagic fever was significantly higher (=0.005, odds ratio=2.5) in those infected with DENV2 (54.8%) when compared to DENV1(32.9%), even though similar proportions of patients had a secondary dengue infection. Patients with DENV2 infection developed leakage significantly earlier (p<0.0001, median= 3, days) when compared to those with a DENV1 infection (median 5 days) and were more likely to develop significant bleeding. Whole genome sequencing showed that these DENV-2 isolates belonged to cosmopolitan strain and was genetically more distant to the DENV-2 strains that circulated from 1981 to 2004 in Sri Lanka.

## Introduction

Dengue viral infections are currently one of the most important vector borne diseases, which is estimated to infect 390 million individuals annually [1]. Although the mortality rates due to dengue have declined in South Asia, the incidence has markedly increased from 285.3 per 100,000 individuals in 1990 to 1371.1 in 2013 and was responsible for 324,200 disability adjusted life years (DALYs) in 2013, which is a sharp increase when compared to 1990[2]. Costs due to hospitalizations and dengue control activities was estimated to be US$ 3.45 million for the year 2012, in Colombo, Sri Lanka [3], which highlights the economic burden due to these diseases in developing, countries.

Although sporadic cases of dengue infection was seen in Sri Lanka in the 1960s with occasional small outbreaks, regular epidemics of dengue only started to occur from year 1989[4]. Since 1989, a marked increase has been seen in the incidence of dengue, with epidemics spreading to many parts of Sri Lanka and during the last few years, the yearly incidence of dengue has been approximately 42,000 cases/year. However, for the year 2017, Sri Lanka reported a total of 186,101 cases of patients who were hospitalized due to dengue, with the yearly incidence increasing from 189.4/100,000 population increasing to 865.9/100,000 population in 2017 [5, 6]. This sudden surge in the number of cases of dengue has resulted in a disastrous situation, where the health care facilities have found it extremely difficult to cope with the high number of patients.

Although the incidence of dengue in Sri Lanka has been steadily increasing, this appears to occur in a step ladder pattern, with a sudden surge in the number of cases every five to six years. In 2009, where there was a sudden 3 fold increase from the previous baseline level, which coincided with the emergence of dengue virus (DENV) serotype 1, which was the predominant serotype until mid-2016 [7]. DENV-2 and DENV-3 were not detected from year 2009 to mid-2016, although the predominant circulating serotypes prior to year 2009, were DENV-2 and DENV3 [7-9]. Since 2009, over 90% of dengue infections were caused by DENV1 and the annual incidence of dengue infection has been around 42,000 cases/year [7, 8]. However, in 2016 the emergence of DENV-2 was detected coinciding with the current large outbreak. Since many unusual clinical manifestations and an increase in disease severity was observed with the emergence of DENV-2, we proceeded to sequence this new strain and to compare the clinical and laboratory features of dengue infections due to this virus, with the previously circulating DENV1.

## Methods

### Patients

172 adult patients with acute dengue infection were recruited from the National Institute of Infectious Diseases of Sri Lanka from year 2015 January to 2017 June, following informed written consent. The day in which the patient first developed fever was considered as day one of illness. Presence of fever, vomiting, abdominal pain, hepatomegaly and bleeding manifestations were recorded daily. Clinical features such as the blood pressure, pulse pressure and the urine output were measured at least 4 times a day. The haematocrit was measured three to four times a day. Serial recordings of full blood count, liver function tests and presence of fluid leakage by ultra sound (US) scan was also assessed daily. Ultrasound scanning was done in the supine position to assess the pleural and peritoneal fluid in the hepatorenal angle, subdiaphragmatic position, in paracolic gutters and in the pelvis.

The severity of acute dengue was classified according to the 2011 WHO dengue guidelines [10]. Accordingly, patients who had a rise in haematocrit above 20% of the baseline haematocrit or detectable fluid in the pleural or abdominal cavities by ultrasound scanning were classified as having DHF. Shock was defined as having cold clammy skin, along with a narrowing of pulse pressure of ≤ 20 mmHg. The day of entering the critical phase was defined as the day from onset of illness, when fluid leakage was detectable by US scan assessment in either the pleural or peritoneal cavities.

### Ethic statement

The study was approved by the Ethical Review Committee of The University of Sri Jayewardenepura. All patients were adults and recruited post written consent.

### Determining viral loads in serial blood samples

RNA was extracted from all serial serum samples using QIAamp Viral RNA Mini Kit (Qiagen, USA) according to the manufacturer’s protocol. The RNA was reverse transcribed into cDNA in the GeneAmp PCR system 9700 using High Capacity cDNA reverse transcription kit (Applied Biosystems, USA) according to the manufacturer’s instructions. Reaction conditions were 10 min at 25°C, 120 min at 37°C, 5 min at 85°C and final hold at 4°C. Multiplex quantitative real-time PCR was performed as previously described using the CDC real time PCR assay for detection of the DENV [11]. This assay was modified to quantify the DENV apart from qualitative analysis. Oligonucleotide primers and a dual labeled probe for DEN 1,2,3 and 4 serotypes were used (Life technologies, India) based on published sequences [11, 12].

### Sequencing of the DEN2 virus

DENV isolation was carried out as previously described using the *A. albopictus* mosquito (C6/36) cell line. The C6/36 cells (approx. 2.5 million cells) were inoculated with 15μl of the RT-PCR-confirmed DENV-2 serum for 1 hour. Cells were returned to fresh media and incubated at 28°C without CO_2_ for up to 12 days or until cells began to show cytopathic effects of infection (displaying non-adherence). Supernatant containing viral particles was harvested and stored at -80°C. Three isolates were initially selected and RNA was extracted from supernatant using SV Total RNA isolation system (Promega, USA), for full genome sequencing. The full genome was amplified for sequencing as 11 overlapping fragments using PCR primers that were created using the Geneious software. The primers used for sequencing is shown in supplementary table 1. Reverse transcription was conducted separately for each fragment similar to work done previously by our group[13]. Sequencing for the above isolate was by the Sanger method by (Macrogen, South Korea) to obtain predominantly dual coverage over the entire genome.

**Table 1:**
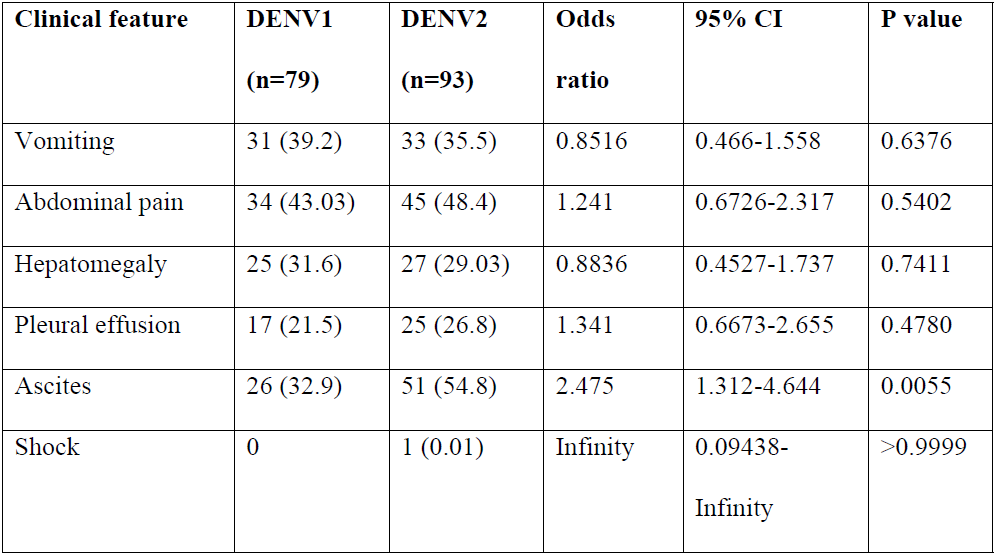
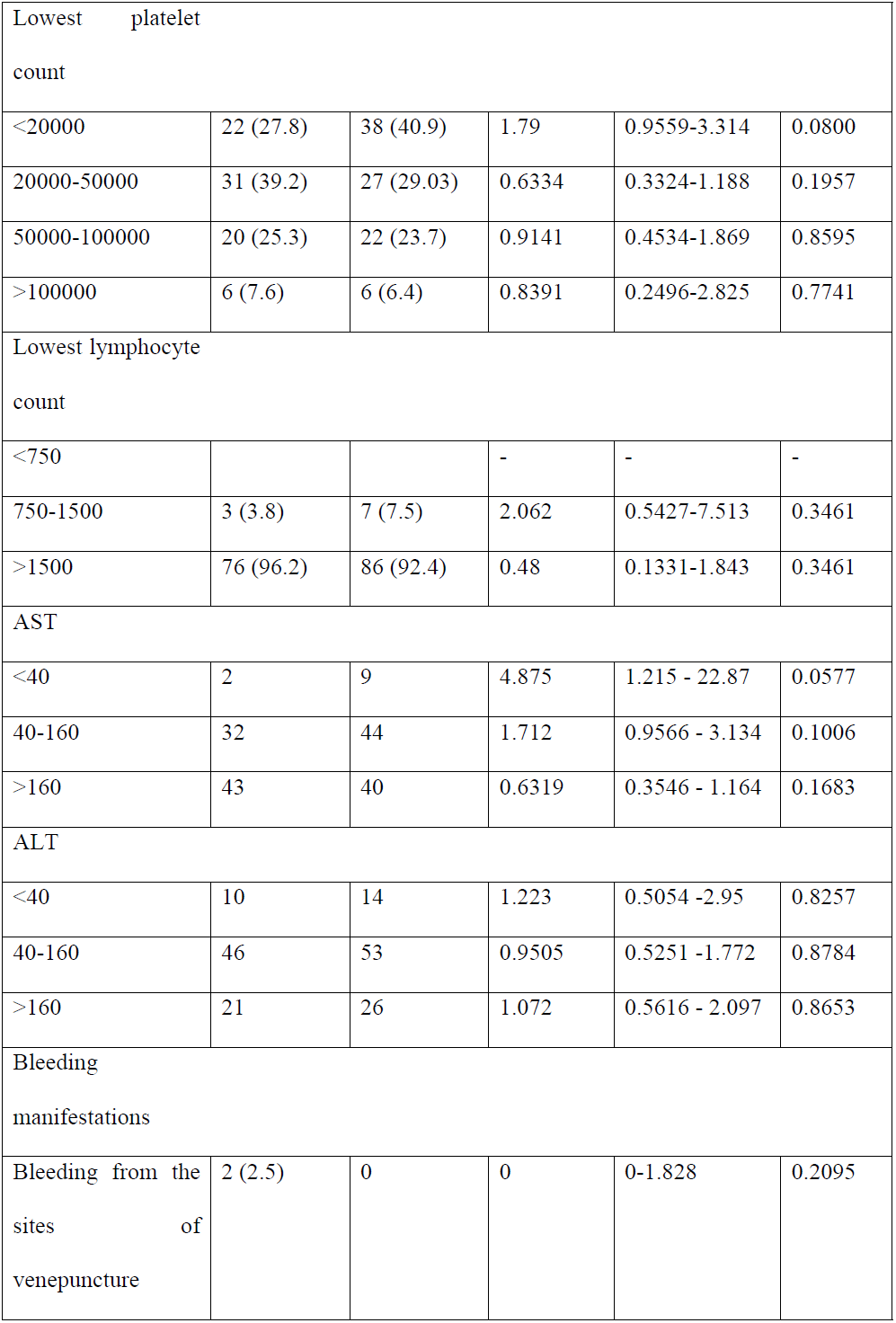
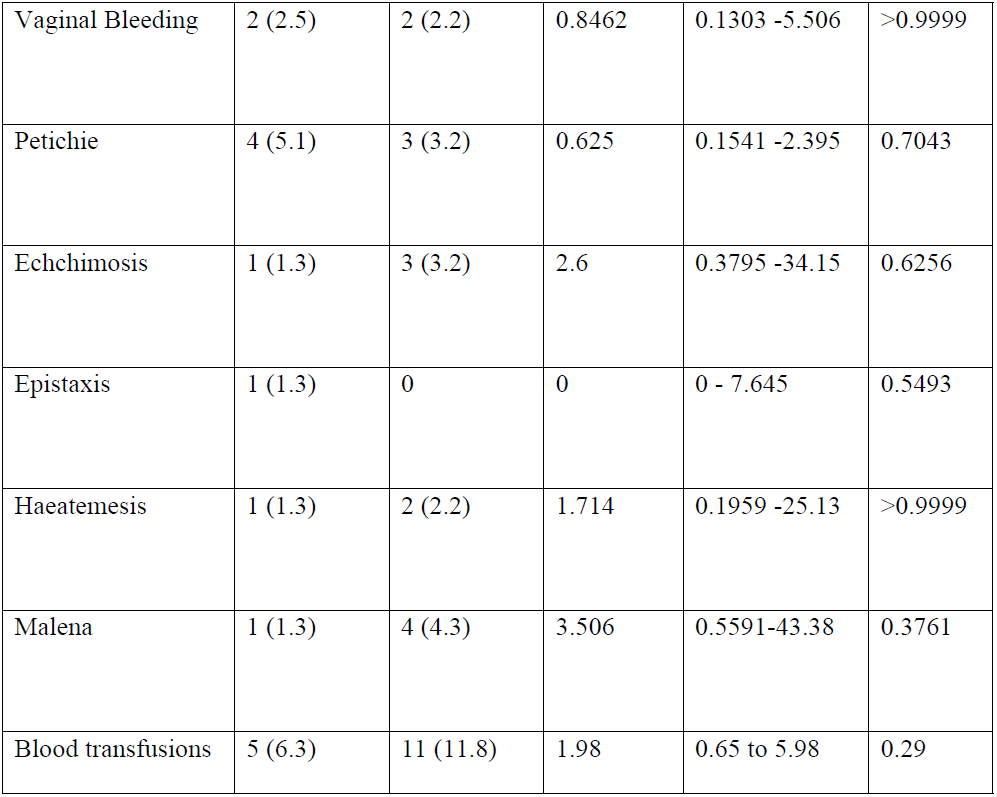
Clinical and laboratory features of with acute dengue infection due to either DENV1 or DENV2.

### Phylogenetic Trees

Sanger sequences were assembled into contigs to generate a full genome as described by Ocwieja et al., 2014[13]. Sequence and identity analysis was performed using GENIOUS software (v10.0.4). Multiple alignments were performed using CLUSTAL W, on GENIOUS, along with phylogenetic analysis using the Neighbour joining method, according to the Tamura-Nei model, with a bootstrap of 1000 replications for analysis of the complete genome and the envelope region of the closely related Sri Lankan strains[14]. Classification and naming of the DENV-2 genotypes were based on the report by Rico-Hesse [15].

### Statistical analysis

Statistical analysis was performed using Graph PRISM version 6. As the data were not normally distributed, differences in means were compared using the Mann-Whitney U test (two tailed). Differences in the serial values of platelet counts and liver enzymes in patients with DHF and DF were done using multiple unpaired t tests. Corrections for multiple comparisons were done using Holm-Sidak method and the statistical significant value was set at 0.05 (alpha). Degree of associations between clinical features due to DENV1 and DENV2 infection, was expressed as the odds ratio (OR), which was obtained from standard contingency table analysis by Haldane’s modification of Woolf’s method. Chi Square tests or the Fisher’s exact test was used to determine the p value.

## Results

Of the 172 patients 79 had an acute DENV1 infection and 93 had an acute DENV2 infection. All the patients with DENV1 were recruited from 2015 to June 2016 and all the patients with a DENV2 infection were those recruited from April 2016 to June 2017. Although few patients had an acute dengue infection due to DENV3 and DENV4, since January 2017 (unpublished data), all patients admitted to this hospital were found to only have DENV2. The details of clinical and laboratory features of these patients are described in table 1. 65 (82.3%) of those with DENV1 and 86 (92.4%) of those with DENV2 were experiencing a secondary infection based on their IgM to IgG antibody ratios. 26 (32.9%) of those with a DEN1 infection and 51 (54.8%) of those with a DENV2 infection developed DHF. Therefore, infection with the DENV2 was associated with a significantly higher risk (p=0.005) of developing DHF (odds ratio=2.5, 95% CI=1.3 to 4.6).

The National Infectious Diseases Hospital of Sri Lanka has a dedicated dengue management unit, where all patients are intensely monitored several times a day. The time of onset of the critical phase where fluid leakage occurs, is defined as when patients develop detectable fluid leakage either in their pleural or peritoneal cavities or when they have a rise in the haematocrit of 20% or more. Patients’ who have a rapid drop in their platelet counts along with a rise in haematocrits and reduction in the urine output, undergo ultra sound scans more frequently to detect fluid leakage. The day of onset of the critical phase is defined as the day from onset of illness, when fluid leakage was detectable by US scan assessment in either the pleural or peritoneal cavities. Patients with DENV2 infection were found to enter the critical phase significantly earlier (p<0.0001, median= 3, IQR 3 to 4 days) when compared to those with a DENV1 infection (median 5, IQR 4 to 5 days).

In addition to those infected with DENV2 were more likely to develop fluid leakage (DHF) and early leakage, they were also more likely to develop significant bleeding in the form of haematemesis and malena (table 1). Only 2 (2.5%) of those with DENV1 developed haematemesis or malena, whereas 6 (6.4%) of those with DENV2 developed such complications. Those with significant overt bleeding or occult bleeding were given blood transfusions according to National Guidelines. Significant occult bleeding was diagnosed clinically if a patient becomes haemodynamically unstable with lowering of haematocrit. While only 5 patients with DENV1 infection needed blood transfusions 11 patients with DENV2 infection needed blood transfusions (p=0.29), indicating clinically significant bleeding is much higher with DENV2, although this was not statistically significant. Furthermore, the proportion of patients who had very low platelet counts (<20,000 cells/mm3) was higher in those who had DEN2 (40.9%) compared to DENV1 (27.8%). Patients with DENV2 had significantly lower platelet counts during day 3, 4 and 5 since onset of illness (Fig 1A). However, there were no differences in the rise in alanine transaminase levels (ALT) or aspartate transaminase levels (AST) during illness in those who were infected with either DENV1 or DENV2 (Fig 1B and 1C). The proportion of those who developed abdominal pain, vomiting and hepatomegaly were also not different in the two groups. Although not significant (p=0.35), the proportion of those who developed leucopenia (<1500 cells/mm3) was higher in those who were infected with DENV2 (7.5%) compared to those with DENV1 (3. %8)

**Figure 1:**
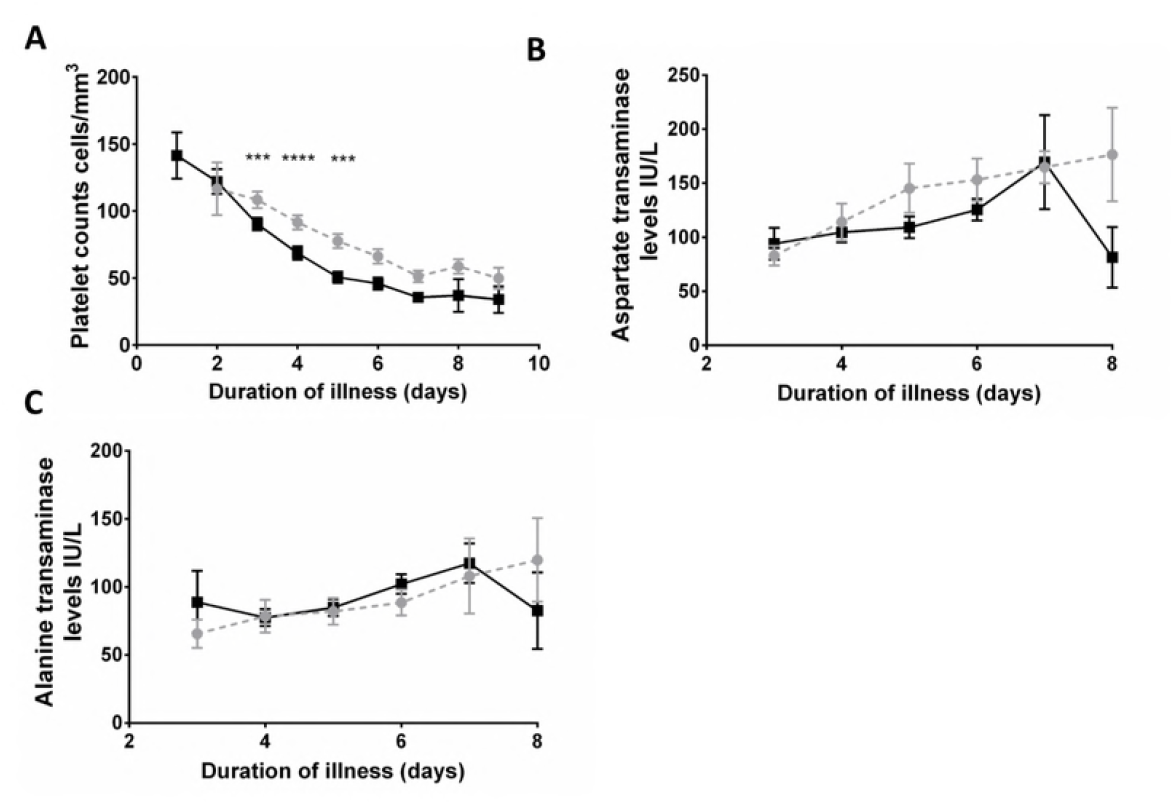
Changes in platelet counts and liver transaminases in patients infected with DENV1 and DENV2. (A) platelet counts of patients were measured daily. The black line represents patients who were infected with DENV2 (n=93), and grey dotted lines represent those infected with DENV1 (n=79). Serum aspartate transaminase levels (B) and alanine transaminase (C) were also measured. Bars represent the mean and SEM *P<0.05, **P<0.01, ***P<0.001, ****P<0.0001.

### Dengue serotype shift resulting in massive epidemic in year 2017

Although the incidence of dengue has been rising in Sri Lanka, the average number of cases reported from year 2009 to 2015 have been approximately, 38,000/year[5]. DENV serotype 1 emerged in year 2009, and this was the only circulating serotype for a few years in Sri Lanka [7, 8]. We have been carrying out surveillance in the National Institute of Infectious Diseases (NIID), Sri Lanka from year 2014 and the changes in the predominant serotypes from January 2016 to January 2017 is shown in figure 2. As shown in the figure, for one year, the predominant serotype, which was DENV1 was completely replaced by DENV2 by January 2017. We have continued to carry out surveillance in this hospital and until December 2017, the predominant and only serotype identified from patients with acute dengue has been DENV2, throughout year 2017.

**Figure 2:**
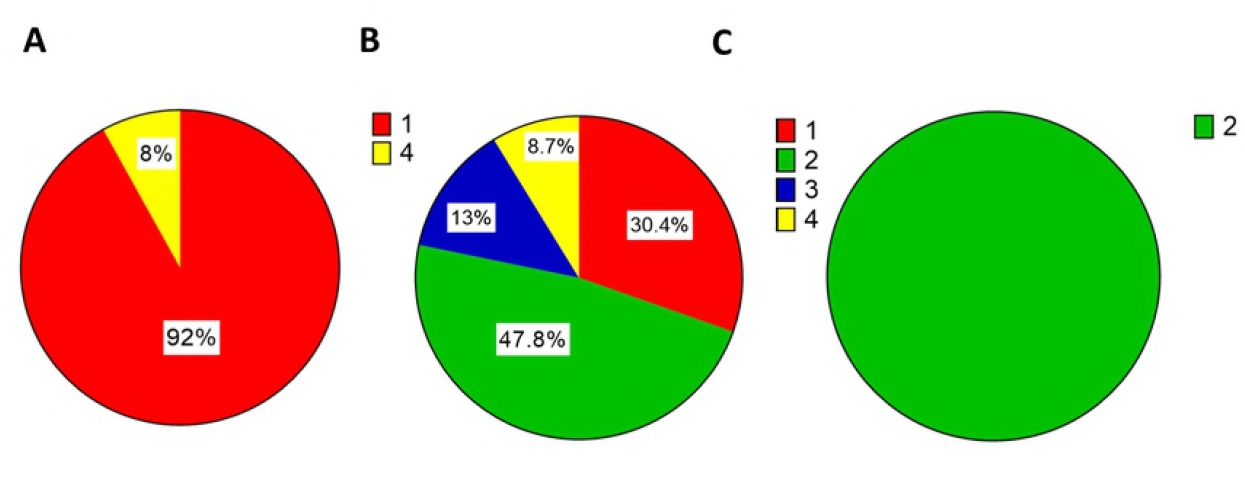
Shift in the dengue virus serotype from DENV1 to DENV2 in year 2016. The DENV serotypes identified from patients who were treated for acute dengue infection at the National Institute of Infectious Diseases during the months of (A) January 2016 (n=75), (B) July 2016 (n=26) and (C) January 2017 (n=48).

### Phylogeography of DENV-2 in Sri Lanka

DENV isolated from samples obtained end 2016 to early 2017 were sequenced. All isolates were confirmed via serotyping PCR as DENV-2 before sequencing. Sequences of 3 Sri Lankan DENV-2 isolates from 2016 were obtained using the Sanger method. We applied the Neighbour joining method according to the Tamura-Nei model to infer the historic spatial dispersion of the virus using our Sri Lankan virus isolates and other reported full length DENV-2 sequences in the literature. We constructed phylogenetic evidence for the course of DENV-2 in Sri Lanka between 1996 and 2016. The earliest recorded full genome sequence from Sri Lanka used in this study was a 1996 strain (FJ882602) and it clusters with the DENV-2 Cosmopolitan genotype viruses (Fig 3A).

**Figure 3:**
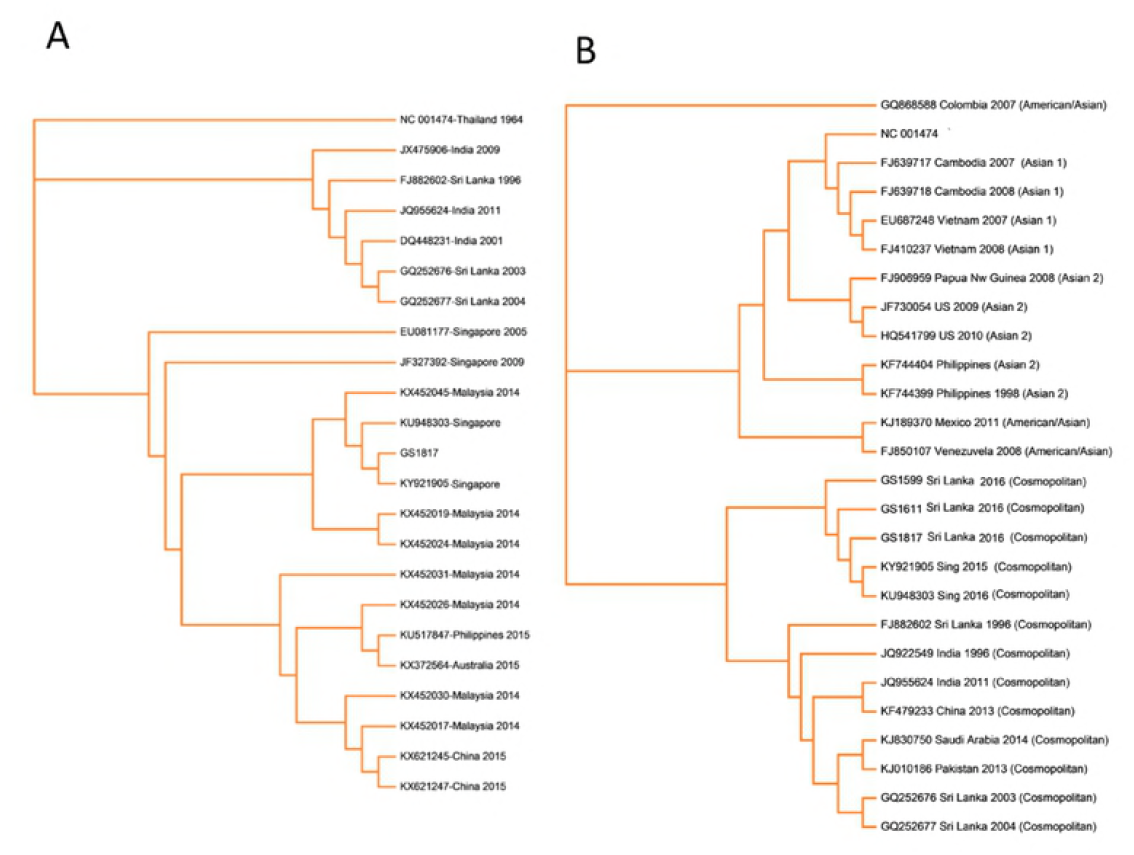
Phylogenetic trees of dengue virus type 2 (DENV-2); A) The tree is based on Full genome sequence of DENV-2 and B) The tree is based on a 239-bp fragment for positions 2311–2550 coding for amino acids at the envelope protein/non-structural protein 1 junction (E/NS1 junction).

The 2016 DENV-2 isolates appear to be closely related to the 2015-2016 Singaporean strains with 99-100% sequence similarity based on the envelope region and the full genome analysis. GS1811 isolate’s full-length genome was compared against several closely related strains. The phylogenetic tree revealed that GS1811 was very closely related to a 2015 Singaporean strain (KY921905), and these two shared a common ancestor with the more recent 2016 Singaporean strain (KU948303).

### E sequence of 2012 Sri Lankan DENV-2 strains

Although the full genome sequencing characterizes all sites of variation in the genome, enabling precise analysis, phylogenetic analyses of DENV traditionally relied on segments of the viral genome, such as the E gene [13]. If the virus travelled to Sri Lanka by way of an additional intermediate country where full-length sequencing is not performed, we would be unable to characterise this by only using whole genome sequencing. Therefore, to improve our resolution we narrowed our database queries to the E recovering the 100 sequences that are most similar to the 2016 Sri Lankan consensus. We used these sequences and other selected DENV-2 sequences to generate a second phylogenetic tree.

The E based analysis confirmed that the 2016 viral strain belonged to the same clade as those strains that circulated in 2015-2016 in Singapore and these shared a common ancestor with the 2014 Malaysian strains. It also confirmed that the more recent DENV-2 strains in Sri Lanka are genetically more distant from the DENV2 strains that circulated from year 1981 to 2004 in Sri Lanka.

## Discussion

Although regular epidemics of dengue infection have occurred in Sri Lanka for over three decades, the largest ever outbreak occurred in year 2017 with the yearly incidence increasing from 189.4/100,000 population increasing to 865.9/100,000 population in 2017 [5, 6]. Results of this study show a complete switch of the dengue serotype to DENV2 in 2017 from predominant serotype DENV1 in 2015/16. This is likely to be the main reason for the large outbreak that occurred recently in Sri Lanka. The incidence rose from 2422 (in May) and 4731 (in June) in the year 2016 to 15,936 (in May 2017) and 25,319 (in June 2017) with an average number of cases increased by more than 3.5-fold. Similar unprecedented outbreaks were reported in Singapore and Malaysia in 2013 with switching of predominant serotype to DENV1 from DENV2 (in Singapore) and to DENV2 from DENV3 and DENV4 (in Malaysia)[16]. Although large outbreaks have been reported in the past from shifting of the DENV serotype from DENV1 to DENV2, the increase in disease severity was due to occurrence of a secondary dengue infection due to DENV2, in a population which had primary infection due to DENV1[17, 18]. In our study, almost equal proportion of those with DENV1 (82/3%) an DENV2 (92.4%) had a secondary dengue infection. Therefore, the increase in proportion of those with DENV2 developing DHF, is unlikely to be due to those with DENV2, having a secondary dengue, while those with DENV1 having a primary dengue infection.

Interestingly, the incidence of dengue infectious appears to increase three-fold every four to six years. Although we do not have data of the shifts in serotypes during the very early epidemics, the sudden rise of dengue from approximately 9500 cases/year from year 2004 to 2009 to 35,000 cases in year 2009, was due to the emergence of DENV1[7, 19, 20]. In year 2017, we again observed a more than threefold rise in the incidence of dengue, from approximately 38,000 cases/year from year 2009 to 2015 to 186,101 cases in year 2017 (Fig 4)[5]. Patients were recruited for this study, was from the Centre for Dengue Management, NIID. The number of admissions for the year 2015 to this unit was 2907 (29.4% of total cases in the Colombo district) for the whole year, whereas for 2017 alone 17,090 patients were admitted (52% of all reported cases in the Colombo district). Therefore, although we only carried out DENV surveillance at the NIID, it is likely to represent the circulation of DENV serotypes in the Colombo district.

**Figure 4:**
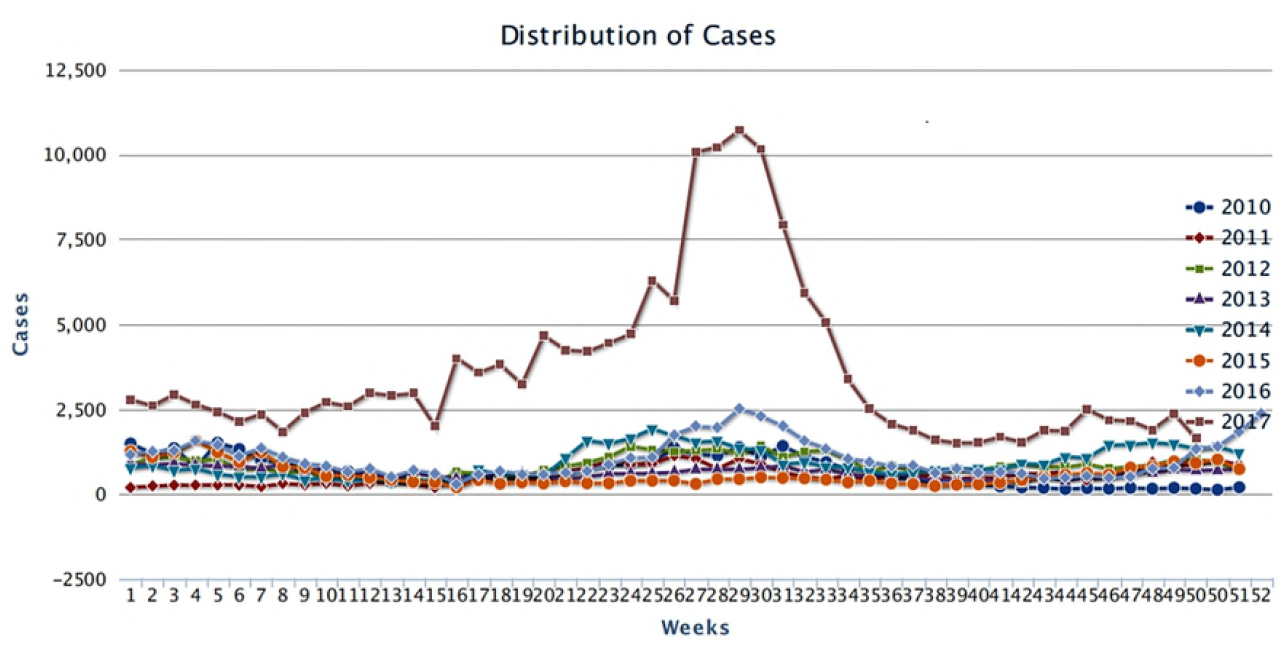
Distribution of reported dengue fever cases, Sri Lanka, 2010–2017. Figure generated using data from the Epidemiology Unit, Ministry of Health, Sri Lanka

During this epidemic, apart from the increase in the number of patients that were admitted to hospital, they appeared to have more complications as well, as 54.8% of patients with DENV2 vs 32.9% with DENV1, developed DHF. 11.8% of patients with DENV2 infection needed blood transfusions compared to 6.3% of patients infected with DENV1. Most importantly, they also developed vascular leakage earlier (median day 3 for DENV2 vs median day 5 for DENV1). During previous years, the advice of the government was to seek medical advice on day 3 of illness as leakage occurred on an average of day 5. However, as patients with DENV2 developed vascular leak and thus developed DHF on an average on day 3 of illness, they were advised to seek medical attention much earlier, further burdening the health care systems[21]. In addition, although the earlier recommendations was to admit patients when platelet counts were below 100,000mm3, along with the presence of dengue warning signs or features such as abdominal pain, persisting vomiting, clinical signs of fluid leakage and mucosal bleeding [22], the admission criteria had to be changed to include patients with platelet counts of ≤130,000mm^3^ [21]. A similar situation, of increase in clinical disease severity was reported among paediatric patients with acute dengue in Thailand, where a significant increase of DHF was reported with DENV2 infection [23]. However, they had not observed an early drop of platelet count, earlier leakage and increased incidence of bleeding, which we have found in our cohort.

As this strain of DENV2 appeared to cause more severe clinical disease, we carried out whole genome sequencing of the virus and carried out analysis of the variations in the envelop gene. We found that current DENV-2 isolates appear to be closely related to the 2015-2016 Singaporean strains with 99-100% sequence similarity based on the envelope region and the full genome analysis. This data is similar to the analysis of the DENV2 sequences isolated from two patients returning to Japan from Sri Lanka this year [24]. A study which was carried out to determine the circulating serotypes in Colombo during the years 2003 to 2006 showed that DENV-2 was seen to be present within the country up to 2006, along with DENV-1, 3 and 4[9]. Kanakaratne et al, showed that DENV-2 strains from 1981 to 2004 were all observed to fall in to the same genotype as the Indian subcontinent/Malaysian (Cosmopolitan) genotype[9]. The study also concluded that there was no evidence of a recent introduction of DENV-2 strain from outside the country due to the close genetic relationship with the DENV2 strains that circulated from 1981 to 2004. The 2015 and 2016 Singaporean and Sri Lankan strains all share a common ancestor with several closely related Malaysian strains from 2014. Although the Sri Lankan 1996 strain and the more recent 2016 strains all belong to the same genotype, the phylogenetic analysis clearly indicates that these two strains are genetically distant, suggesting that the more recent strains entered the country from Singapore (based on the reported data). The same differences were observed between the 2016 strains and the 2003-2004 strains from Sri Lanka. The 2004-2003 strain clustered with the 1996 strain and shared a more common ancestor with each other while the more recent strains clustered together sharing another common ancestor. However, it is interesting to note here that during the 2009-2014 DENV-1 epidemics there were no DENV-2 infections reported in Sri Lanka and until recently it did not make an appearance.

In summary, the largest ever dengue outbreak of Sri Lanka, which occurred in 2017 appears to be due to the shift in the circulating serotype from DENV1 to DENV2. This virus was also more likely to cause DHF and significant bleeding. In addition, the fluid leakage developed significantly earlier than due to other serotypes. Since this DENV2 strain appears to cause more severe forms of clinical disease, it would be important to determine variations in the virus genome or other immunological factors that could have possibly contributed to severe disease.

